# Transformation of event representations along middle temporal gyrus

**DOI:** 10.1101/468199

**Authors:** Anna Leshinskaya, Sharon L. Thompson-Schill

## Abstract

When learning about events through visual experience, one must not only identify which events are visually similar, but also retrieve those events’ associates—which may be visually dissimilar—and recognize when different events have similar predictive relations. How are these demands balanced? To address this question, we taught participants the predictive structures among four events, which appeared in four different sequences, each cued by a distinct object. In each, one event (‘cause’) was predictably followed by another (‘effect’). Sequences in the same relational category had similar predictive structure, while across categories, the effect and cause events were reversed. Using fMRI data, we measured *associative coding*, indicated by correlated responses between effect and cause events; *perceptual coding*, indicated by correlated responses to visually similar events; and *relational category coding*, indicated by correlated responses to objects in the same relational category. All three models characterized responses within right middle temporal gyrus (MTG), but in different ways: perceptual and associative coding diverged along the posterior to anterior axis, while relational categories emerged anteriorly in tandem with associative coding. Thus, along the posterior-anterior axis of MTG, the representation of the visual attributes of events is transformed to a representation of both specific and generalizable relational attributes.

## INTRODUCTION

The typical predictive relations among events form an essential component of world knowledge, even with respect to everyday objects. To understand what it is for something to be a *light switch*, one must understand the effect of flipping it; to be a *poison*, the effect of ingesting it; and to be a *plant*, the importance of watering it. It is the way these objects participate in contingencies per se that is essential, because to say that plants often get watered, or sometimes wilt, does not capture the essentially contingent property that the plant will wilt *if* it is not watered (Goldwater and Gentner 2015; Mumford 1998; Jones and Love 2007; Gentner 1983; Pinker 1989). Here we investigate the cognitive and neural mechanisms supporting long-term memory of predictive relations, a relatively neglected topic in cognitive neuroscience.

There are two challenges that such representations present to the cognitive system. First, specific predictive relations can hold among events which look nothing alike—such as flipping a light switch and a lamp turning on—making them often at odds with visual similarity (Hindy, Ng, and Turk-Browne 2016; Kok and Turk-Browne 2018). It thus poses a distinct demand from discriminating visually distinct stimuli. One would want to both relate a switch-flipping and a lamp turning on, as well as recognize that two different lamps turning on are similar events, different from switch-flipping events—whether or not they are related predictively. As we review below, both functions have been related to the ventral temporal lobes, creating a puzzle regarding how both of these functions can be simultaneously accomplished when they are in conflict.

The second challenge is that predictive knowledge itself must be generalizable to be a useful component of semantic memory. We need to not only represent that *this* switch turns on the light, but recognize the functional similarity between different switches, buttons, and knobs that also control lights. This requires recognizing not just the specific relation between one switch and one lamp, but use analogy to recognize similar relations across different circumstances: an ability termed relational category formation (Jones and Love 2007; Kemp et al. 2010; Gopnik and Meltzoff 1997; Markman and Stilwell 2001; Goldstone, Medin, and Gentner 1991; Markman and Gentner 1993; Christie and Gentner 2010; Corral and Jones 2014; Stuhlmueller, Tenenbaum, and Goodman 2010). Importantly, not all switches that look alike share this relational property—some might control ceiling fans, be ineffectual, and so on. Our understanding of the neural mechanisms of relational category representation are extremely sparse, as does our understanding of how we build them from event experience.

The present experiment aims to address these puzzles by measuring all three kinds of representations simultaneously: visual discrimination of events, long term memory of predictive relations between specific pairs of events, and relational categories grouping similar specific relations across contexts into a common class. We hypothesize that cognitively, relational categories could be built by relying on specific predictive representations, and thus, we anticipate a close relationship between neural representations of them. On the other hand, we expect both predictive representations to pull apart from those of visual similarity, as these functions are at odds.

Prior work offers an elegant way to probe the neural basis of long-term memory of specific predictive relations, by examining which areas show correlated responses to individual presentations of visual stimuli after learning their association (e.g., Sakai & Miyashita, 1991). Only after learning, visually responsive neurons previously tuned specifically to stimulus A increase their response to associated stimulus B, even when A and B are no longer presented together. This signature captures an important part of what it means to represent a (specific) relation: given that perceptual similarities between associated stimuli are controlled, the only reliable commonality between associated pairs is the fact of their association. Thus, a common response between them would seem to represent this fact. We use the term “associative coding” to designate this signature.

Much neurophysiological research has found associative coding signatures in higher-level ventral visual stream areas, specifically anterior-medial aspects of macaque inferior temporal (IT) cortex (Higuchi and Miyashita 1996; Erickson and Desimone 1999; Miyashita 1988; Naya, Yoshida, and Miyashita 2003; Messinger et al. 2001; Sakai and Miyashita 1991). These areas span macaque area TE, an apex of the ventral visual stream, and perirhinal and entorhinal cortices, more associated with memory. In partial accordance with this work, human fMRI has found evidence of associative coding in ventral stream areas like PPA and FFA (Turk-Browne et al. 2010; Senoussi et al. 2016; Favila, Chanales, and Kuhl 2016; Polyn et al. 2005; Zeithamova, Dominick, and Preston 2012) but also in earlier visual areas such as V1 (Hindy, Ng, and Turk-Browne 2016)^1^.

As we noted earlier, if predictive relations are represented in areas responsible for visual discrimination, it creates something of a puzzle. How could ventral stream areas simultaneously distinguish faces from houses (e.g., Haxby, Gobbini, & Furey, 2001), and represent an associated face and house similarly? Surely, these functions would interfere with each other. On this basis, one might speculate that predictive representations are more strongly represented outside of the specific areas that encode visual similarity and support discrimination of those particular events from each other (what we term “perceptual coding”). However, past work has not directly compared these functions.

Nonetheless, in line with this intuition, associative coding has also been found outside of ventral stream areas, notably in the hippocampus, to a sometimes stronger or fuller degree (Schapiro, Kustner, and Turk-Browne 2012; Hindy, Ng, and Turk-Browne 2016; Kok and Turk-Browne 2018). For example, Hindy et al (2016) found that hippocampal representations capture more of the full sequence of a set of events than visual prediction in V1, and Kok et al (2018) found that V1 responses are dominated by an on-screen stimulus more than what is predicted from it. However, these findings regarding V1 does not address the rest of the ventral stream. Furthermore, hippocampal responses may be limited to recently learned, pre-consolidated predictive knowledge. Others have reported that ventral stream areas broadly conceived are not be the strongest ones to represent predictive content, and find different cortical areas that do (Long, Lee, and Kuhl 2016; Kuhl and Chun 2014). Overall, both sets of findings bolster our prediction that associative coding and perceptual coding diverge neurally. We test this idea directly by creating orthogonal models of these forms of coding and directly comparing their signatures.

There is much less prior work regarding the second puzzle: whether neural representations of specific predictive relations generalize to similar relations among visually different stimuli in distinct contexts (the signature of relational categories). Only two experiments, to our knowledge, have investigated such relational categories in the brain. Frankland and Greene (2015) probed agent vs patient roles as expressed syntactically in language; they examined where sentences like “the truck hit the ball” elicited similar neural responses to sentences like “the ball was hit by the truck” (same relation), but different from “truck was hit by the ball” (different relation but similar surface features). They found this pattern specifically in left lateral superior temporal cortex (near superior temporal gyrus), as well as information representing which object participated in which role. A somewhat similar paradigm found a nearby area, among others, but did not do inferential testing of cortical location (Wang et al. 2016). In neither case is it clear that similar results would obtain for memory representations, as opposed to syntactic analysis. We thus performed a whole-brain search for relational category information as retrieved from memory of events, with the aim of understanding how they relate to specific predictive representations (associative coding).

To do this, we taught participants predictive relations among four events by presenting them in continuous sequences (Figure 1). There were four distinct sequences, each involving a similar set of events, but appearing with or around a differently-shaped object. Each sequence contained a strongly predictive pair of events, which we call the *cause* and the *effect* (rather than a cue and an outcome, as there were no prespecified “cues” in our paradigm)^2^. Between sequences/objects, we varied which particular events served as the cause and the effect, so as to create relational categories among them. In one category, the objects were “causers”: their movements preceded the relevant ambient event (following the example in Figure 1, the light flash). In the other, the objects were “reactors”: the light flash was now the cause, and the objects moved in response to it. The two objects in the same category always exhibited different movements (whole-body tiling vs moving a detachable part) to ensure that surface similarity went against the grain of the relational categories. Participants were then scanned about a week following learning, to ensure we probed consolidated, long-term memories. During the scan, we did not show any sequence information, but rather, had participants retrieve it from memory as they viewed the individual events in random order alongside the objects.

**Figure 1.**
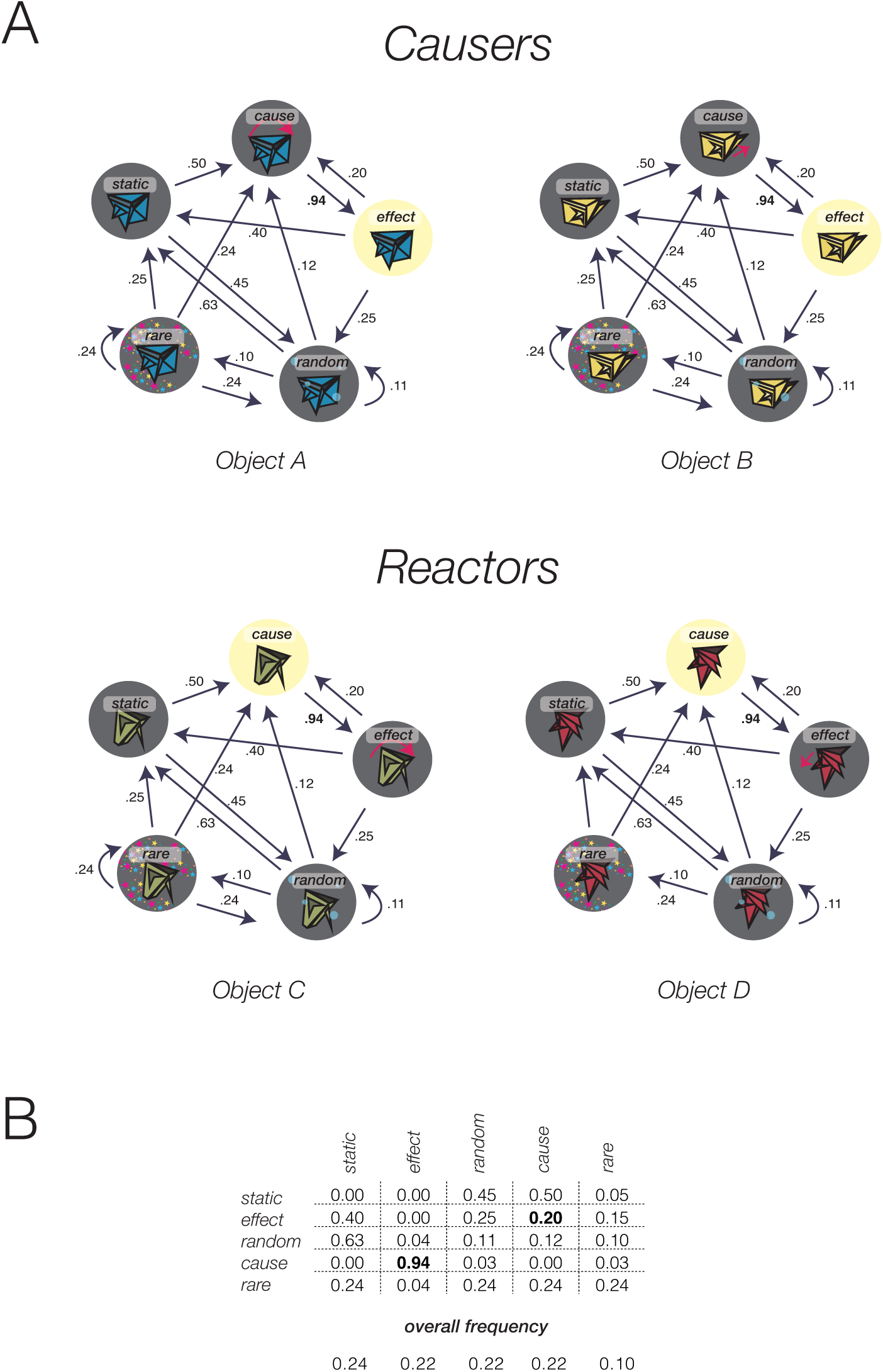
**A**. The four object shapes used to cue each training sequence (counterbalanced) and transition probability structure among the events within those sequences. The relations among the abstract roles we the same for all sequences, but event assignments differed. In two of these, object movements precede a specific ambient event (*causers*); in the other two, object movements follow the same ambient event (*reactors*). The identity of the ambient event serving as cause/effect was counterbalanced across participants. **B**. The transition probability structure among all event roles in each sequence represented in matrix form; below, overall frequency of each event.

To probe specific predictive relations/associative coding, we measured the extent to which the individual cause and effect events elicited correlated neural patterns, relative to weakly predictive pairs, within the context of the same object. To probe relational categories, we compared events in the context of different objects to each other, testing whether there is a broad similarity between seeing events with object A (which tilts followed by light flash) and object B (which moves a detachable part followed by light flash), relative to object C (which tilts before a light flash). This kind of representation thus captures a generalized relation: the relation that a light flash is predictable, even across visually different causes (part move vs tilting), and across distinct contexts, but different from a light flash being the predictor of similar movement events (tilt and part-move).

Finally, to probe visual similarity/discrimination (“perceptual coding”), we measured the similarity of neural responses to the same event vs different events as seen in the context of different objects–for example, whether a light flash surrounding object A was similar to a light flash surrounding object B, relative to a movement of object B. We then tested how the cortical locations of these three forms of representation were related to each other.

## METHOD

### Participants

Participants were recruited from the University of Pennsylvania community via the Experiments@Penn website. Procedures were approved by the Institutional Review Board at the University of Pennsylvania and all participants provided written informed consent.

Participants in Session 1 were required to be between 18-35 years old, with no history of neurological disorder, and right-handed. To continue to Session 2, they had to be eligible for MRI following detailed screening, and achieve > 80% performance on a forced choice test at the end of the Session 1 learning task. This, rather than lengthening the training, was done to keep constant the amount of training exposure across participants. Two participants served as a pilot sample for parameter testing and were not included in reported analyses. 97 additional participants performed Session 1. Of these, nine did not meet MRI eligibility criteria during the detailed screening; one was excluded due to participating in a related prior experiment; ten were unable to be scheduled for Session 2 within the targeted time-window; and 24 were excluded based on their performance on the training task. This exclusion rate is relatively high, but follows the performance cut-off specified in the pre-registration (see Procedure). 53 participants took part in Session 2 and underwent fMRI. Of these, a total of 17 were excluded due to the following reasons: substantial misunderstanding of the task (1), failure to form relational categories as assessed on a post-scan measure (3), technical glitch causing data loss (1), completion of fewer than 8 runs due to delays or discomfort (6), and excessive motion (6). The final sample included 36 participants (25 female), with a mean age of 23 (range 18 - 35).

### Registration

The methods of this experiment were pre-registered at https://osf.io/3mj4v/. Major deviations from the registration are noted in the manuscript, and minor ones on a document available at this URL. Most notably, we increased our sample size from 24 to 36 following two major unexpected outcomes: 1) inability to find associative coding in MTL as expected based on prior literature, and 2) inability to find any region at the whole-brain corrected level representing relational categories. The latter prevented us from being able to test our hypothesis about where such representations would be localized. However, following the addition of 12 participants, these facts of our data did not change; thus, the sample size increase is not likely to have inflated the significance level of analyses we do report. Inclusion criteria were as pre-specified, but we additionally required that participants showed evidence of having formed relational categories, as this would otherwise hamper our ability to find such representations in cortex. Several analyses were not pre-planned in the pre-registration or had minor deviations. This includes the perceptual coding analysis, which was originally planned to be cross-object (grouping the two objects which tilted vs. moved a detachable part); we later realized the analysis would be better matched to the associative coding analysis by analyzing individual events. It also includes the vector-of-ROI analysis, which was designed to follow up on our significant searchlight findings.

### Overview of session structure

Participants completed two sessions, which were 3–11 days apart (*M* = 6)^3^. Session 1 took about 2 hours, and involved a training task (see Procedures). At Session 2, participants reviewed what they had learned in Session 1, then underwent fMRI scanning while performing a retrieval task, and answered a post-scan questionnaire.

### Stimuli

Stimuli are illustrated in Figure 1. They consisted of four novel geometrical objects, each embedded 5 distinct types of animated events, presented as GIFs: bubbles, stars, light flashes, movement (either whole-body tilt or local part movement), and static (object still on screen). Each event was composed of twelve 100ms frames (total duration 1200ms), except the static event (total duration 2400ms). Frames were hand-drawn using Adobe Illustrator and concatenated into GIF files using Matlab (Mathworks).

For the training task, these event stimuli were concatenated into 450-event-long sequences, one for each object separately; this created the “object contexts”. The order of events in each sequence followed a specific structure, as summarized in the pairwise transition matrix shown in Figure 1B (and in graphic form in 1A). This matrix specifies the conditional probability of moving into any specific state at any trial n given the state at trial n-1. Sequences for each participant was generated probabilistically using a weighted walk, where the probabilities of adding events to the sequence were specified by this transition matrix. We ensured that the generated sequences closely matched this specified probability structure by checking that, in each generated sequence, the average absolute difference in all pairwise transition probabilities was below .00004, and the standard deviation of state frequencies of the cause, effect, and random were below 4. The actual average obtained transition matrix was nearly identical to the specified sequence.

Although the same transition matrix governed the abstract structure of the predictive relations in all four sequences, the way that the events were assigned to this structure varied (Figure 1A). In all cases, a strong predictive relation held between two events, the *cause* and the *effect*, such that the cause is followed by the effect with a 94% probability. For two objects, the *causers*, the cause was the object’s movement (tilt for object A, and part-move for object B), and the effect was one of the 3 ambient events (bubbles, stars, or light), selected for each participant in counterbalanced fashion, but always the same for the two objects (e.g., light flash in the example in Figure 1C&D). For the other two objects, the effect and cause events were swapped: object C tilted following the ambient event (e.g., light flash), while object D moved a detachable part following the same ambient event (also light flash in this example). In this way, object contexts belonged to one of two relational categories, causers and reactors.

Two other events served as *random* and *rare* events, which were almost never predicted by the cause, and almost never predicted the effect; the identity of these events was the same across categories. The random event was matched in frequency to the effect and cause, while the rare event was half as frequent; overall frequencies are displayed in Figure 1B. This difference in frequencies was introduced to enable comparison to a planned follow up, in which categories were based on frequency rather than contingency, but is not a manipulation of interest here. Comparisons largely involve the cause, effect, and random events.

The assignment of object shapes to relational category was counterbalanced across participants, creating 6 counterbalancing conditions for object shape (i.e., all possible assignments to two categories). Relational category was orthogonal to object movement, as the two members of each category always had different movements. The assignment of the 3 event ambient types (light flash, bubbles, and stars) to be the *effect/cause*, *random*, and *rare* was also counterbalanced across participants, creating 6 other counterbalancing conditions, which were paired randomly with the shape counterbalancing conditions.

During fMRI, participants saw individual event gifs for each of the 4 non-static events (cause, effect, random, and rare) in the context of each of the 4 objects, creating 16 conditions. As described in Procedure, this presentation was in random order, rather than following the transition probabilities as during training.

### Procedure

#### Session 1

In Session 1, participants were introduced to the four object shapes and told that their task was to learn which events are likely to follow which others, in the presence of each object. The 450-event sequence for each object was split into a preview block and 3 task blocks. The preview showed the first 50 events (~1 minute) from each object’s sequence, with the order of sequences/objects random. The following 3 blocks showed the remaining 400 events from each sequence. These were presented in sets by block number, such that participants saw block 1 for all 4 objects, then block 2 for all 4, and so on, with the order of sequences/objects randomized uniquely at each block.

In these task blocks, the videos were interspersed with intermittent questions, which probed the participant to decide what event is most likely to come next (Figure 2A). The response options showed static images of two other events (not itself or rare), and a “both equally” option. For example, following the presentation of a cause event, participants would choose between the effect, random, or “both equally”. The “both equally” option was correct for events with close or equal transition probabilities (within 5 percent), but otherwise the correct option was defined as whichever event had the higher conditional probability. For example, following the cause, the right answer was always the effect. However, following the effect, cause and random were equally likely; and following random, cause was more likely. Presentation side of the two event options was randomized, with “both equally” appearing below. Participants received feedback following their response, showing them which response was correct if they were incorrect. For each object, there were 10 questions pertaining to the cause, effect, and random events, and 6 pertaining to rare, creating 36 questions total, distributed randomly over the 400 events. To create the 3 blocks, the sequences were split so that each one contained 12 questions for each object (hence these segments could vary in the number of events).

**Figure 2.**
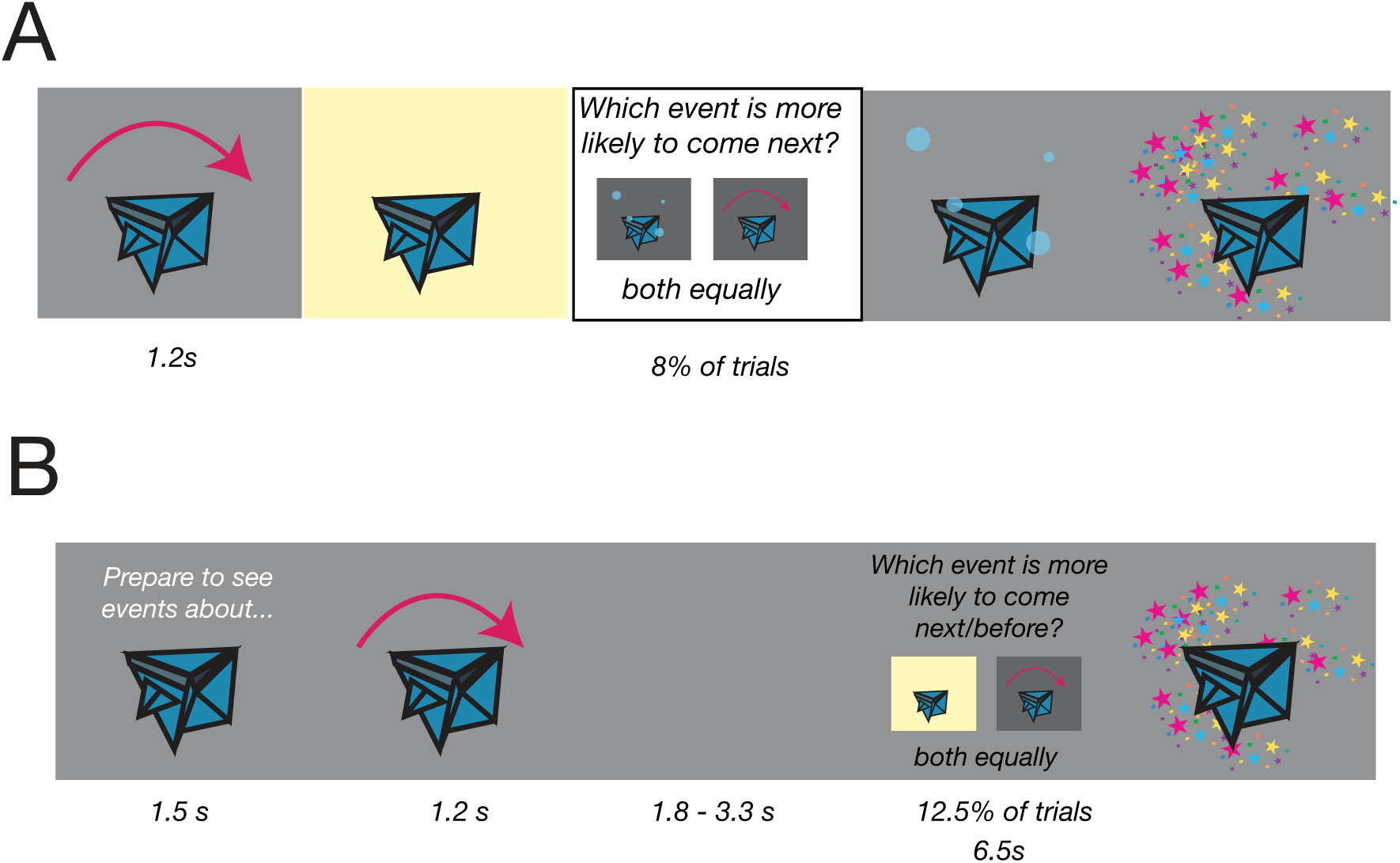
**A**. Example of the training task; sequence presentation is interspersed with questions probing what will come next. **B**. Example of the in-scan retrieval task. Each block began with a cue, followed by 4 events involving the same object interspersed with delay periods. 12.5% of events were followed by a question probing either what would typically come next, or before, in the previously learned sequences.

Following each block, participants saw forced-choice tests probing their knowledge about each object’s sequence. Each question showed two videos side by side, where each video contained a sequence of two events (e.g., tilt followed by light flash vs. tilt followed by bubbles). Participants had to choose which of the two videos was most typical. There were 8 trials per object, including 3 trials comparing cause–effect to effect–cause, 1 trial comparing cause–effect to random–effect, 1 trial for cause–effect to random–cause, and two filler trials (effect–cause vs random–effect, and effect–cause vs random–cause), which balanced the number of times the cause–effect pair and the effect—cause pair were shown overall. Participants had to obtain 80% or higher on the non-filler questions by block 3 to continue to Session 2. The emphasis on cause-effect order knowledge was because of their importance for the relational categories.

#### Session 2

In Session 2, participants reviewed what they learned with a similar training task as in Session 1, but shorter: a total of 18 questions was shown per object over 240 events, split over 2 blocks. During fMRI scanning, participants performed a retrieval task in which they saw individual events separated by blank screens, and in randomized order except blocked by object. They were intermittently asked questions probing their memory of typical event order from training. Figure 2B illustrates the retrieval task. Each block began with a cue (1.5 s) showing a still image of the object, followed by 4 of that object’s events (these could be any of the 4 events in any order, but with no more than 2 of the same event in a row). Each event (1.2 s) was followed by a delay (blank screen) with a duration of 1.8, 2.3, 2.8, or 3.3 s. On 12.5% of trials, this delay was followed by a question (6.5 s). Half of the questions asked what is likely to come next; the other half asked what is likely to come before. This was so that participants could not anticipate making any specific response. Each question had three response options, as in the training task: two options were images of two other events (except itself or rare), and one was “both equally”. The presentation side of the two event images was randomized. Unlike the training task, there no feedback was given, except an overall score at the end of each run. Order of object blocks was randomized.

Each of the 16 events (cause, effect, random, and rare for each of the 4 objects) was shown exactly 4 times in each run, once with each delay duration. These were arranged randomly into 4-event blocks (with the constraint that no event could repeat more than twice in a row within a block). There were 10 runs over the entire experiment, and thus 40 repetitions of each event. Questions were distributed across the entire 10 runs, ensuring that there were 5 questions for each event (12.5%), and half were ‘after’ and half were ‘before’ questions. For analysis of fMRI data, question periods were modeled separately and not further analyzed.

Following scanning, participants completed a questionnaire asking whether they thought the objects could be naturally grouped into categories (yes/no), and, if so, how many and on what basis (freeform text entry). They were then shown draggable images of each object shape and asked to arrange them on the screen such that the ones they thought were most similar were closer together and the ones they thought were most different were furthest apart. Finally, they answered questions about their perceptions of causality and animacy. The animacy question asked, “To what extent did the 4 objects you learned about seem like animate, living objects (animals/people) vs. inanimate (non-living objects like artifacts)?”, with a 1 – 5 response scale where the endpoints were labeled ‘Definitely Animate’ and ‘Definitely Inanimate’, with the side of the scale of these labels randomized. Another question probed their perception of causality regarding the “causer” objects: “For two of the objects, their movements predicted the occurrence of another event (e.g., the appearance of a light flash, bubbles, or stars). To what extent did you perceive this relationship as causal? Did these objects seem to cause this event?”, and showed a similar 1 – 5 response scale whose end points were labeled Definitely Causal and Definitely Not Causal, and the ends of the scale randomized. A third question probed causality about the reactor objects (“Did the event seem to cause the object to move?”).

### fMRI acquisition parameters

fMRI data were acquired using a Siemens Magnetom Prisma 3T scanner at the University of Pennsylvania, using a 64-channel coil. Anatomical volumes were acquired with a T1-weighted MPRAGE sequence with 0.8×0.8×0.8mm voxel resolution, 256mm field of view, TR = 2.40s, and TE = 2.24ms. Functional data were acquired with a multi-band EPI BOLD sequence using 72 interleaved slices with a multi-band acceleration factor of 3, 2×2×2mm in-plane voxel resolution, 220mm field of view, TR = 2.0s, TE = 30ms, and flip angle = 75°. Slices were aligned to the posterior-anterior axis of the hippocampus (following Schapiro et al., 2012).

### fMRI preprocessing

Data were pre-processed using AFNI software (Cox 1996). Slices in each volume were corrected for acquisition timing using Fourier interpolation (3dTshift). Each volume was spatially aligned to the 4^th^ volume of the first scan to correct for motion (3dVolReg). Image intensities were normalized (scaled to range from 0 – 100) and linear and polynomial slow trends up to the 3^rd^ level were removed. Data were spatially smoothed using a Gaussian kernel of 3mm full-width half-maximum. Runs in which displacement from the first exceeded 3mm were excluded; if more than two were excluded, the dataset was discarded for that participant (and replaced to fulfill the counterbalancing set). Included runs were then concatenated into one time-series and entered into linear modeling. Anatomical volumes were spatially aligned to the first functional volume in line with the rest of the functional data, then spatially transformed to Talairach space. Parameters for Talairach transformation were then applied to functional scans for volume analyses.

### Linear modeling

Two linear models were fit to the data. The Object model included regressors for each object presentation (A – D), which spanned cue periods, event trials, and delay periods, but excluded question periods, which were modeled with a separate regressor. Derivatives of the 6 motion realignment parameters (4 directions and 2 rotations) were also included. The Event model included regressors for each of the 16 different events (cause, effect, random, and rare for each of the four objects), and, as above, a regressor for all question periods and 6 motion realignment parameter derivatives. In both models, volumes with motion outliers (those with > .15mm displacement from the previous) were excluded.

Regressors were created by convolving the time-courses of each condition in each run with a gamma-shaped hemodynamic response function. The convolved time-courses were then used as predictors in a least-squares linear regression over the time-courses of BOLD signal in each voxel (3dDeconvolve). This produced a map of regression coefficients for each condition, and their respective t-values, reflecting the slope of the relationship between that voxel’s signal and the occurrence of that condition. The t-value maps were used in all subsequent analyses.

### Anatomical surface analysis

Anatomical volumes were converted to surface maps for surface-based searchlight analyses. Surfaces were created using the Freesurfer function recon-all (Fischl et al. 1999), which used intensity gradients to segregate white and gray matter and generate inflated cortical surface maps for each individual participant. This algorithm also performed segmentations of medial temporal lobe areas which were used in ROI definition (see below). Inter-individual alignment of surface maps, and alignment of functional data to surface maps, were performed using AFNI (mapIcosohedron) and algorithms implemented in the Surfing toolbox (Oosterhof, Wiestler, and Diedrichsen 2010; Oosterhof et al. 2011).

### MVPA analyses

We defined three voxel-wise similarity models for multi-voxel pattern analyses. These models reflected different predictions about which conditions in either the Event or Object linear models should be relatively more vs. less similar (correlated) in terms their voxel-wise responses (i.e., t-values in a set of voxels in a searchlight or ROI region). The models are depicted in Figure 3. For each analysis, pairwise correlations among the conditions of interest (in terms of voxel-wise Tests of model fit against 0 were performed with one-tailed t-tests because a model fit below zero is not meaningful, as greater neural similarity between conditions or stimuli which are *different* on the dimensions of interest is not interpretable. We describe each of the models in turn below.

**Figure 3.**
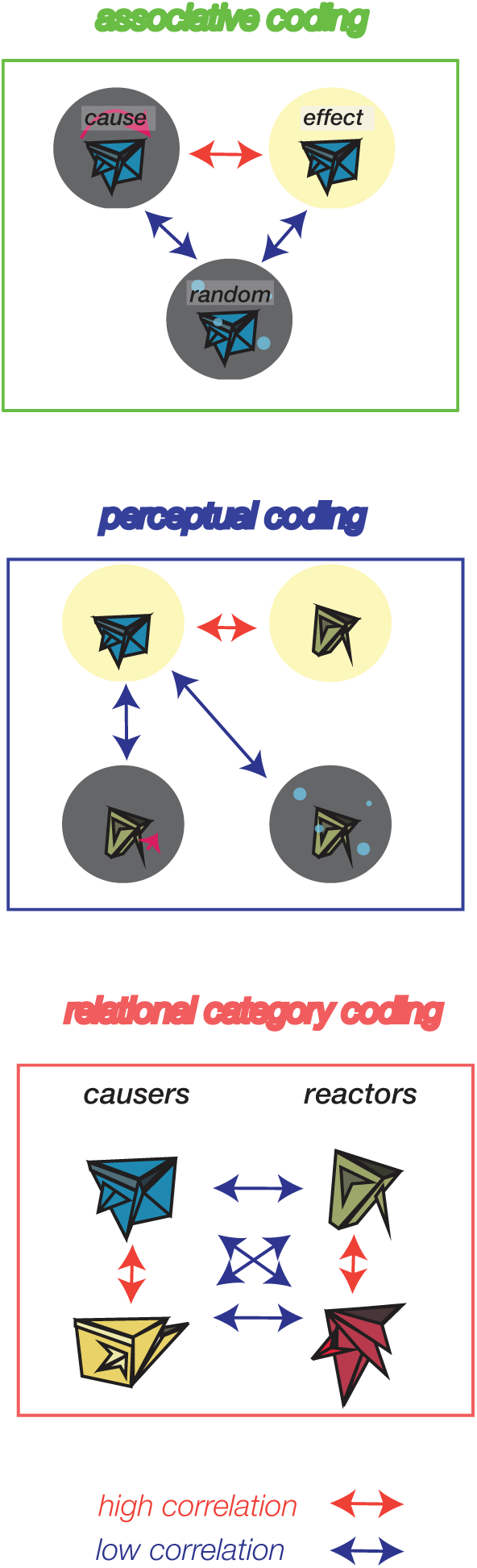
Illustrations of the three models used to fit the neural data. Associative coding predicts that events which reliably predicted each other should more elicit similar voxelwise responses than those which occurred in the context of the same object but did not reliable follow each other. Perceptual coding predicts that stimuli which are visually similar should elicit similar neural patterns relative to those which are visually dissimilar (as shown in the context of different objects). Relational coding predicts that objects cueing structurally similar sequences (e.g., the two causers) elicit similar neural activity.

#### Associative coding

Following prior work (Schapiro, Kustner, and Turk-Browne 2012), we looked for representations of specific predictive relations by comparing the voxel-wise responses to strongly predictive events (*cause* and *effect*) relative to weakly predictive events (*cause* and *random* and *effect* and *random*), within object context. These comparisons were chosen because the stimuli assigned to be effect and random were counterbalanced across participants, thus fully controlling for stimulus identity. We subtracted these two correlation values within each object, and then averaged across objects to obtain an overall measure of associative coding in a region, which we refer to as the ‘associative coding model fit’.

#### Perceptual coding

Viewing a light flash in the context of object A is perceptually more similar to a light flash in the context of object B than bubbles in the context of object B. We queried where such perceptual similarity might be represented by comparing the average pairwise correlation of similar events across objects (when these were either effect, cause, or random, but not rare, because the latter is not matched in frequency), relative to the correlation between different events across objects (e.g., the correlation between the light flash with object A and the light flash with object B, vs. the light flash with object A and movement, bubbles, or stars with object B). This average correlation difference was computed for each participant in each region and is referred to as the ‘perceptual coding model fit’.

#### Relational category coding

In a generalized representation of a relation, sequences in which the light flash is an effect should be more similar to each other than sequences in which light flash is a cause. Accordingly, our 4 object/sequence conditions could be grouped into two classes based on similar relational structure, that is, whether the object’s movement predicted vs. followed one of the ambient events (causers vs reactors). We used the outputs of the Object linear model to correlate the voxel-wise t-values between all pairs of objects, and subtracted the correlation of all different-relation objects (e.g., object A and object D) from the correlation of same-relation objects (objects A & B, and objects C & D). The average difference was computed for each subject in each region as is referred to as the ‘relational category model fit’.

### Anatomical ROI definition

Anatomical ROIs tested were left and right hippocampus, entorhinal and perirhinal cortices, as well as individual hippocampal subfields (CA1, CA3, CA4, dentate gyrus, subiculum, and tail), all as extracted from the Freesurfer segmentations of each individual. We also tested a left middle superior temporal region based on coordinates reported in Frankland & Greene (2015), at (−59, −25, 6), by defining spherical ROI with 123 voxels surrounding them. MVPA analyses as described below were performed on the voxels within each ROI in each participant.

### Searchlight analyses

Searchlight analyses enabled us to test our three models across cortex by defining neighborhoods of contiguous voxels tiling the brain. Searchlights were defined both on volume and surface data, with the latter preferred as it defines searchlight neighborhoods respecting the curvature of individuals cortical surfaces, a more valid measure of contiguity (Oosterhof et al. 2010; Oosterhof et al. 2011). We report clusters seen in both analyses, but performed multiple comparison correction only on surface maps due to computational time constraints. Volume searchlights were used for reporting Talairach coordinates for clusters significant in the surface analysis. Each searchlight neighborhood had a radius of 3 voxels or 6mm and included 123 voxels. An additional follow-up analysis targeting medial temporal areas used a 3mm radius. All pairwise correlations among conditions in their voxelwise t-values were computed, fisher-corrected, and subtracted and averaged according to the respective model, yielding a single value for that neighborhood reflecting model fit. Subsequently, t-tests were used to compute the statistical significance of model fit at each searchlight neighborhood across the group. Surface maps were created to display the value for each neighborhood at its center coordinate.

Multiple comparison correction was performed with permutation testing at the cluster level. For each individual, 10 null maps for each linear model were created by shuffling the condition labels across trials (i.e., randomizing trial-to-condition assignments). Then, for each of 1000 permutations, one of these null maps was chosen per participant at random, and analyses proceeded exactly as they had been for real data. The maximal cluster size obtained from group-level analyses at each iteration was used to build a distribution of maximal cluster size expected by pure chance (noise), given a pre-cluster threshold of *p* < 001. The observed cluster sizes in the real data were assigned a significance value based on their likelihood in this distribution; clusters above 105 mm^2^ were significant at *p*_corr_ < .05.

#### Functional ROI definition

We used the significant clusters from the whole-brain, surface searchlight analyses with the associative coding and perceptual models to define individual functional ROIs. To do so, we used the group clusters as boundaries, then selected individual surface nodes in each participant by taking the largest cluster of non-zero nodes within it. These regions were then tested for fit of the other models, all of which used independent comparisons (between different trials) from those used to define the ROIs. This enabled us to test the theoretical question of whether the same or different regions enable associative and relational coding.

#### Vector of ROI analysis

To statistically assess the spatial relationships between perceptual and associative coding results, which we found in middle temporal gyrus (MTG) at a whole-brain-corrected significance level, we took a “vector of ROIs” approach (Konkle and Caramazza 2013). To do so, we defined a linear axis along MTG, respecting its boundaries along the surface curvature. We then defined spherical ROIs along this axis, taking every 10^th^ node and defining a sphere around it with a radius of 10mm; this ended up creating 39 partially overlapping ROIs along the posterior to anterior axis of MTG. We extracted the fit of the perceptual, associative, and relational category models to assess their relationships across these ROIs.

## RESULTS

### Learning Performance – Session 1

Accuracy on questions interspersed through the training sequences was high among the included subjects (*M* = 82%, *SD* = 13%). For two participants, training task responses were missing due to technical glitches. Accuracy did not differ as a function of whether the object was a causer or a reactor (*t*(33) = 1.04, *p* = .304). Accuracy did differ as a function of the event probed, with the causal event being most accurate (*M* = 86%, *SD* = 13%), followed by the random event (*M* = 78%, *SD* = 16%), the effect event (*M* = 72%, *SD =* 25%), and the rare event (*M* = 68%, *SD =* 22%). These differences were significant for all comparisons between cause and others, and between random and effect (Table S1). Such differences no doubt arose because the predictive relations from the cause were by far the strongest and clearest, while the predictive relations among the other events were weaker. It should be noted that the associative coding model predicts relatively higher correlations between cause and effect than between cause and random (or between random and effect); in terms of accuracy, however, the effect event was most different from the cause event in terms of accuracy. Furthermore, the correlation across subjects between accuracy on the cause and effect events was lower (*r* = .60) than between the cause and the random event (*r* = .70) and between the effect and the random event (*r = .*78). Therefore, the difficulty of the training task itself is not confounded with the neural models tested (in fact, it goes in the opposite direction).

Forced-choice tests probed the ability to retrieve each cause-effect relation associated with each object, and was used as a selection criterion for scanning. Included participants were thus highly accurate on each object, reaching ceiling by the last block (*object 1, M =* 0.98%, *SD* = .08%; object 2, *M* = 98%, *SD* = 8%; object 3, *M* = 98%, *SD* = 8%; object 4, *M* = 98%, *SD* = 8%), with no difference between causers and reactors (*t*s < 1) either at the last block or on average across all blocks.

### Learning Performance – Session 2

In Session 2, participants performed a review task similar to the Session 1 training. Accuracy remained high (*M* = 85%, *SD* = 13%) and followed a similar pattern across event types as in Session 1. Participants again reached ceiling on the forced-choice test (*M* = 98%) with no difference between object categories. During the in-scan task, participants were also highly accurate (*M* = 74%, *SD* = 16%). Accuracies for questions about the cause (*M* = 67%, *SD* = 19%) were relatively different from accuracies about the effect (*M* = 81%, *SD* = 14%) as well as about random (*M* = 80%, *SD* = 16%). The correlation between participants’ accuracies were not significantly higher between cause and effect (*r = .*64) than between cause and random (*r =* .52, *z*(35) = 0.73*, p* = .459) or between random and effect (*r* = .37, *z*(35) = 1.50, *p* = .134).

In a post-scan measure, participants were asked to spatially arrange images of the four objects according to their judgments of how similar they were to each other. Four participants’ data were missing due to technical error. We computed the screen distance between all pairs of objects. Participants reliably placed the same-category objects (the two causers and the two reactors) closer to each other than to different-category objects (*M* = 50.33, *t*(31) = 12.00, CI [58.88 41.78], *p* < .001). All but one considered them to belong to two categories, rather than any other number, despite no suggestion in the experiment that they should do so. Thus, the included participants spontaneously grouped the objects into two categories according to the predictive structure of their associated sequences.

Participants reliably saw the objects as inanimate, providing a mean rating below the midpoint of 3 on a 1—5 animacy scale (*M* = 2.31, SE = 0.22, *t*(35) = −3.25, *p* < .01), though there was a wide range on this measure (1—5), indicating that some participants did see the objects as animate. They also rated the object movements as reliably *causing* the ambient event for the causers (*M* = 4.04, SE = 0.20, *t*(35) = 5.27, *p* < .001) and the ambient event causing the object movement for the reactors (*M* = 3.93, SE = 0.21, *t*(35) = 4.53, *p* < .001), with no difference between the two (*t*(35) = 1.00, *p* = 0.32). It should be noted that no causal language was used at any point during the experiment.

### Areas Exhibiting Associative, Perceptual, and Relational Category Coding

We performed whole-brain searchlights to localize neighborhoods of voxels showing the characteristic ‘associative coding’ signature of specific predictive representations: higher correlation among pairs of events which were predictive during learning (cause and effect), relative to pairs of events which were not predictive of each other (cause and random, and effect and random), within each the context of each object. We found such representations in a diverse set of regions across cortex, as shown in Figure 4A; this included right MTG and lateral prefrontal cortex, left precuneus, and medial prefrontal cortex. Some of these resemble parts of the default mode network (Buckner, Andrews-Hanna, and Schacter 2008). However, we did not find predictive representations in medial temporal or inferior temporal regions. Volume-based searchlights broadly confirmed these findings (Figure S1); Talairach coordinates are reported in Table S2. In anatomically defined medial temporal ROIs (hippocampal, parahippocampal, perirhinal and entorhinal cortices), we found no significant effects (all *t*s < 1, except in the right parahippocampal gyrus, in which *M* = 0.013, *t*[35] = 1.32, *p* = .098). We additionally tried a smaller searchlight radius, but again found no effects at *p* < .01 uncorrected anywhere in the medial temporal lobes apart from a small cluster in left parahippocampal gyrus. Thus, overall, we found little evidence of associative coding in MTL, but robust evidence in other areas.

**Figure 4.**
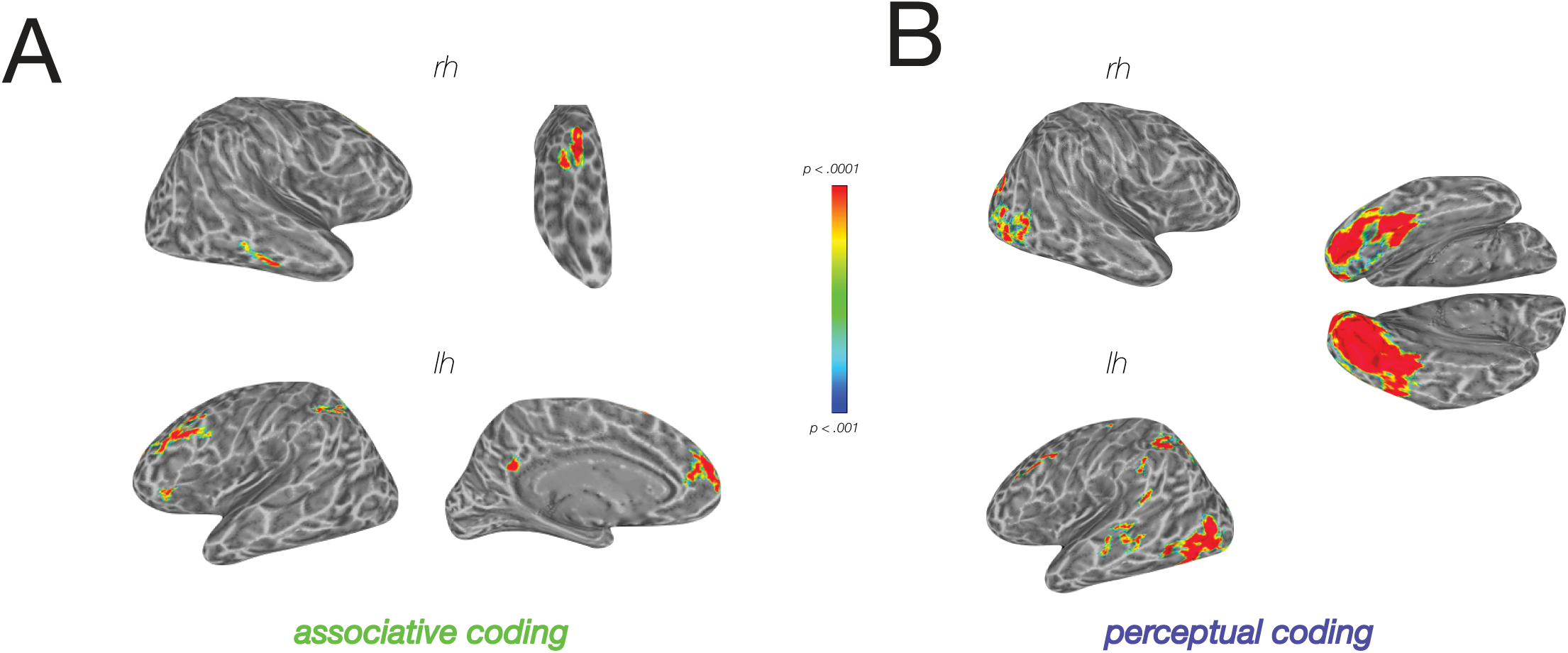
**A**. Surface-based searchlight results for associative coding, with a voxelwise threshold of *p* < .001 and a cluster-corrected threshold of *p* < .05 (104 mm^2^). **B**. Surface-based searchlight results for perceptual coding, with the same thresholds as A.

We did not find any region at the whole-brain level reliably exhibiting relational category coding, i.e., greater correlation when viewing objects that cued sequences with similar statistics (the same effect and cause events), relative to objects with different statistics. We also failed to find effects in an ROI centered on coordinates reported in Frankland & Greene (2015), *M* = 0.004, *t*(35) = 1.07, *p* = .147. This may be due to lack of power, or to the specificity of these prior effects to language stimuli or their extraction from syntax.

Lastly, we searched for regions that contained information about the perceptual qualities of the stimuli—those which showed correlated activity between similar events across objects (e.g., light flash in Objects A and light flash in Objects B, C and D) relative to visually dissimilar events across objects (light flash in the context of Object A and movement of Objects B, C, and D). We found such effects across occipito-temporal cortex, spanning medial and lateral IT (Figure 4B). Lateral areas of IT are known to be particularly sensitive to dynamic events, with MTG showing selective responses to inanimate motion (Beauchamp et al. 2002). These previously reported coordinates (−46, −70, −4, Talairach space) are very near the peak of our perceptual coding effects in lateral temporal cortex (47, −71, 3). Notably, although both perceptual coding and associative coding engaged parts of right MTG, these areas seemed non-overlapping.

### Posterior to Anterior Functional Divisions in MTG

To further validate this spatial divergence statistically, we performed a vector of ROIs analysis in individual participants (Konkle and Caramazza 2013). We defined a series of partially overlapping ROIs along the posterior-anterior axis of right MTG in order to directly test that the spatial relationship between associative and perceptual coding is statistically reliable, and assess the relationship between these model fits to those of the relational category model.

Figure 5A shows these ROIs, and the fit of each model (associative, perceptual, and relational category, in terms of correlation difference) in each ROI, arranged from posterior to anterior. This reveals that perceptual coding declines, while associative and relational category coding increases, along this axis. This is statistically reliable: the location of the peak ROI for the associative model (*M* = 19.81, *SE* = 1.53, CI [16.7386, 22.8725]) is reliably anterior to the peak of the perceptual model (*M* = 12.44, *SE* = 1.69, CI [9.0586, 15.8303]) when compared in individual subjects (*t*(35) = −3.51, *p* = 0.001, *d* = −0.76). Fitting linear slopes to individual data along the ROIs confirms that associative coding exhibits an overall positive slope (*M* = 0.001, *SE* = 0.0005, CI [0.0000, 0.0021], *t*(35) = 2.06, *p* = .047) while perceptual coding exhibits a negative slope (*M* = −0.001, SE = 0.0003, CI [-0.0017, −0.0005], *t*(35) = −3.85, *p* < .001) and that these are significantly different from each other (*t*(35) = 3.75, *p* < .001, *d* = 0.87). These analyses confirm that a divergence between perceptual and associative coding along MTG holds reliably when comparing their locations in individual participants (something which is not guaranteed by the results of the searchlights).

**Figure 5.**
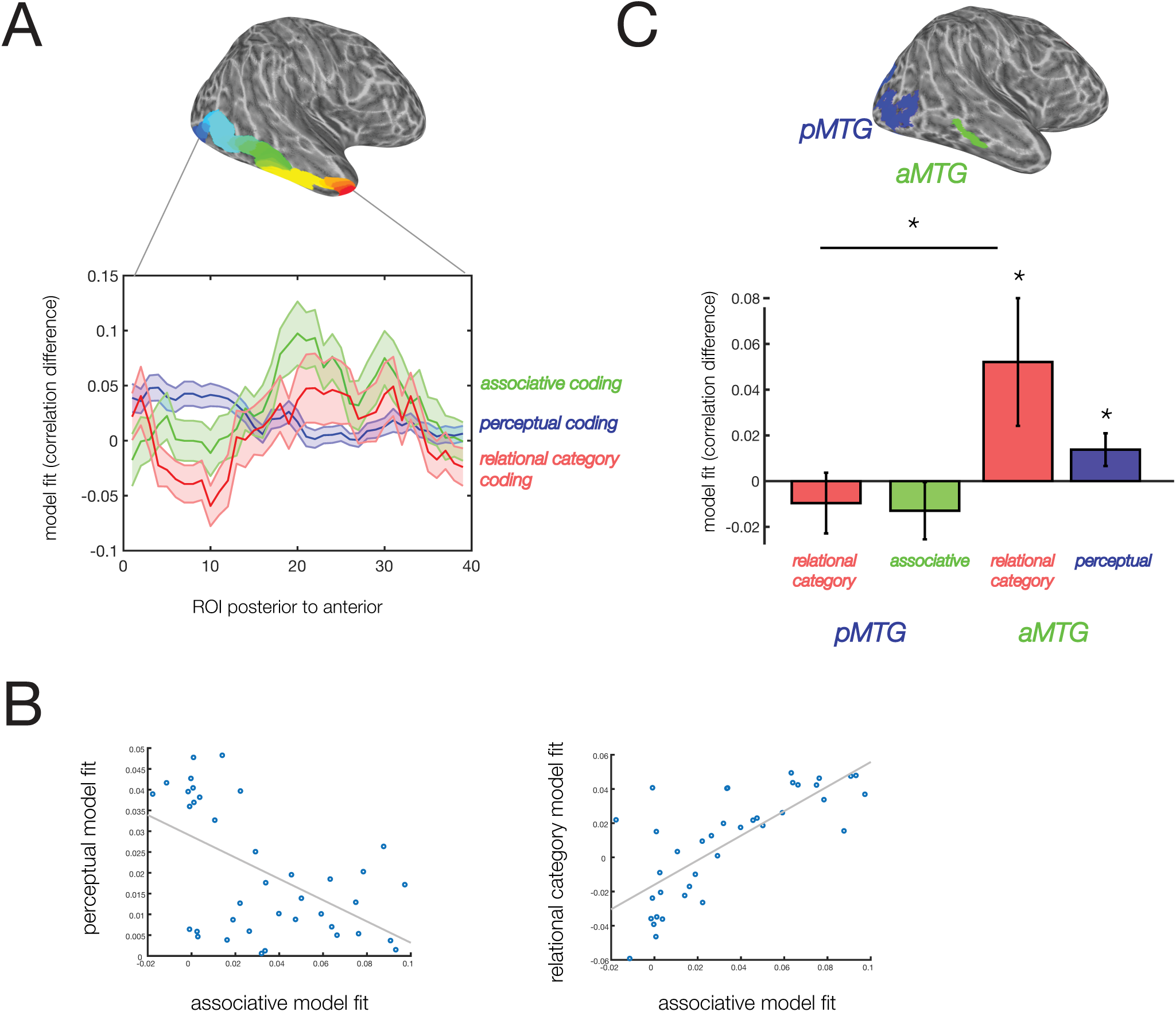
**A**. Results of the vector-of-ROIs analysis along the posterior-anterior axis of right MTG. ROIs, color coded by number, are shown above; below, the fit of each of the three models (associative, perceptual, and relational category) is plotted against the index of each ROI. **B**. Correlations between the associative and perceptual models (left) and associative and relational category models (right) across the mean fit values in each of the 39 ROIs in MTG. **C**. Functional ROIs based on the associative model searchlight (aMTG) and perceptual model searchlight (pMTG), shown above, and the fit of the relational category model, associative model, and perceptual model, excluding the model used to define each ROI, below. * = *p* < .05, uncorrected for multiple ROIs.

We additionally explored how associative and perceptual coding relate to relational category coding across these ROIs. It is visible in Figure 5A that the fit of the relational category model along right MTG (red line) emerges in tandem (i.e., spatially co-varying) with the associative model fit (green line), while the perceptual model fit (blue line) diverges from both. This is statistically evident in a peak location analysis: the location of individual participants’ relational category model peak (*M* = 19.81, *SE* = 2.05, CI [15.6966, 23.9145]) is on average the same as the location of their associative model peak (*p* = 1), but is anteriorly shifted from the perceptual model peak (*t*(35) = −3.13, *p* = 0.004, *d* = −0.65). The slope of the relational category model fit is a positive but not significantly different from 0 (*M* = 0.0009, *SE* = 0.0006, CI [-0.0003, 0.0020], *t*(35) = 1.54, *p* = .134), but it is significantly more positive than that of the perceptual model (*t*(35) = −3.50, *p* = .001). We further assessed this relationship by computing the correlation between the three models in terms of their fit (averaged across participants) along the 39 ROIs; these correlations are shown in Figure 5B. The correlation between perceptual model fit and associative model fit was strongly negative, *r*(37) = −0.54, *p* < .001, while the correlation between relational category model fit and associative model fit was strongly positive, *r*(37) = 0.74, *p* < .001, with a significant difference between them, *M* = −1.29. z(37) = −6.36, *p* < .001. We performed a complementary test within individual participants, by taking each individual’s vector of model fits across the 39 ROIs, computing the correlation among models, and testing these individual, fisher-corrected correlation values against 0 at the group level. Here we found a negative but non-significant correlation between associative and perceptual model fits, *M* = - 0.009, *SE* = 0.0580, *t*(35) < 1, but a significant positive correlation between associative and relational category model fits, *M* = 0.095, *SE* = 0.0459, CI [0.0027, 0.1864], *t*(35) = 2.09, *p* = 0.044, *d* = 0.35, though the difference between these two correlations was not significant, *t*(35) = 1.50, *p* = .144.

Although statistical tests of the associative and perceptual model fits relative to 0 are biased, as their locations informed the ROI definition, the fit of the relational category model is independent. We found that the relational category model fit did not significantly exceed 0 when correcting for the 39 ROIs, but that several anterior ROIs showed effects at uncorrected significance levels (ROI 31: *M* = 0.049, *SE* = 0.027, CI [0.0043, Inf], *t*(35) = 1.85, *p* = .036, *d* = 0.31; ROI 33: *M* = 0.040, *SE* = 0.0213, CI [0.0049, Inf], *t*(35) = 1.92, *p* = .031, *d* = 0.32, one-tailed tests).

A key feature of our design is that participants’ task queried both forwards and backwards predictive relations (what comes next *and* before), so that participants would retrieve memory for both causes and effects in all conditions. However, one might argue that, despite these instructions, participants primarily retrieved forwards predictions. If so, the correlation between relational category and associative coding could be driven by the fact that, in similar relational category blocks, participants predominantly retrieved the same event from memory (the effect, when seeing the cause). If these assumptions are true, then we should not see the same relative correlation effects between our three models when using trials showing the effect, random, and rare effects. We tested this idea. Despite the reduction in the amount of data included, we found a consistent set of results: relational category coding was negatively correlated across ROIs with the perceptual model, *r*(37) = −0.32, *p* = 0.048, but still positively correlated with the associative model, *r*(37) = 0.69, *p* < 0.001. Peak locations along the right MTG significantly diverged between relational category and perceptual coding effects, *M* = 5.69, *t*(35) = 2.26, *p* = 0.03, but did not diverge between relational category and associative coding effects, *M* = −1.67, p = .500. However, we did not see effects in individual subject correlations. Overall, however, this suggests that the correspondence between associative and relational category effects across right MTG is not driven only by the retrieval of a particular event during cause trials, but by the entire predictive pattern across the events in each condition.

In summary, we found evidence that relational category and specific associative coding are related in terms of their location along right MTG, while perceptual coding diverges from both, particularly at the group level (i.e., in terms of which models are reliable across subjects). In individual participants, relational category coding was anterior in peak location to perceptual coding, but overlapping with associative coding, and was correlated in fit across ROIs only with associative coding (though not significantly more so than with perceptual coding). When considering the cross-subject reliability of model fits across ROIs, we found that ROIs with stronger relational coding were more likely to have stronger associative coding, but *less* likely to have strong perceptual coding. Altogether, these data suggest that specific associative coding emerges spatially in tandem with increased relational category coding, but diverges from perceptual coding, along right MTG.

### Functional ROI analysis

Another way to probe these model relationships is to define individual ROIs based on the whole-brain searchlight results for associative and perceptual coding, and examine the fit of the relational category model within them. We defined individual ROIs within the significant group clusters in right anterior right MTG (aMTG), which was of particular interest given the vector of ROI findings, as well as right prefrontal cortex (PFC), left PFC, left intraparietal sulcus (IPS), left medial prefrontal cortex (MPFC) and right precuneus (PC). Results from the perceptual model searchlight were used to define right posterior right MTG (pMTG). The aMTG and pMTG ROIs are shown in Figure 5C.

In aMTG, we found significant fits for the relational category model, *M* = 0.052, SE = 0.027, CI [0.0056, Inf], *t*(35) = 1.89, *p* =.033 one tailed, *d* = 0.32, and the perceptual model, *M* = 0.013, *SE* = 0.007, CI [0.0019, Inf], *t*(35) = 1.96, *p* = .029 one-tailed, *d* = 0.33, with no difference between them, *t*(35) = 1.34, p = 0.189. In pMTG, neither the associative nor the relational category model had significant fits (relational category model, *M* = −0.0096, *SE* = 0.0133, CI [-0.0317, Inf], with t(35) = −0.73, p = 0.766; associative model, *M* = −0.0129, *SE* = 0.0125, CI [-0.0338, Inf], t(35) = −1.05, p = 0.845). Thus, in line with the vector of ROI analysis, the fit of the relational category model was significantly stronger in aMTG than in pMTG, *M* = −0.0096, SE = 0.0133, CI [-0.0317, Inf], *t*(35) = 2.62, *p* = .013, *d* = 0.47, the latter of which was not significant, *t* < 1. Overall, these analyses suggest that aMTG contains information about relational properties, both specific and general, as well as about the perceptual properties of events, while pMTG contains only perceptual information.

It should be noted that the relational category and perceptual model effect in aMTG do not survive correction for the testing of multiple ROIs. However, we had prior reason to test this region given the results in the vector of ROI analysis, which suggested that this is where perceptual and associative coding may most diverge and show differential relationships to relational coding. This analysis confirms those findings, but the significance of the fits themselves should be interpreted with caution.

A different way to consider the relationship among models is to look at the correlation among model fits across participants within these functional ROIs. Here, we did not find that participants with higher associative model fits in aMTG had higher relational category model fits in this area, *r*(34) = −0.05, *p* = .774. Instead, we saw a marginal correlation between associative model and perceptual model fits, *r*(34) = 0.30, *p* =.074, though this correlation was not significantly higher (*z*(34) = 1.46, *p* = .144). We did not see significant correlations among models in pMTG. The contrast between these results and those of the vector-of-ROI analysis suggests that the relationship between models manifests specifically at the level of large-scale organization: that associative and relational category coding are both more likely to be found anteriorly, rather than posteriorly, while perceptual coding is more likely to be found in posteriorly, rather than anteriorly. Within these ROIs, however, model fit relationships across participants do not follow the same pattern, and could simply reflect that participants who paid more attention to the events had higher fits for perceptual and associative models, while relational category models had some independent variance.

In the remaining associative coding ROIs, we found significant fits of the perceptual model in left PC, left MPFC, left LPFC, and left IPS (Table S3). Thus, even though these areas did not have strong enough perceptual model effects to appear in searchlight findings, most associative areas did contain information about the perceptual properties of the events. However, we did not find significant fits of the relational category model in any ROI (all *t* < 1). Thus, the pattern of divergence and overlap that we found in right MTG may be specific to this area.

Finally, we did not find any significant correlations between our behavioral measures (of accuracy or categorization) and model fits in any ROI, perhaps because participants were close to ceiling on these measures by design and thus exhibited limited range.

## DISCUSSION

We investigated the neural mechanisms supporting long-term predictive memory by probing three kinds of representations that the brain might simultaneously exhibit when processing visual events: their visual features (perceptual coding); memory of their specific predictive relations (associative coding); and their generalized relational categories (relational category coding). We aimed to better understand the relationship between these kinds of representations in order to resolve two puzzles: (1) how the brain simultaneously encodes the distinctions between visually distinct events while also representing the predictive relations among them, and (2) how it might build generalizable representations of predictive relations across distinct contexts.

We found a diverse set of areas representing specific predictive relations, but these areas were not the ones showing the strongest evidence of perceptual coding. Perceptual coding was found to be strongest in posterior aspects of the temporal lobe, including posterior MTG, an area known to be important for processing dynamic stimuli (Beauchamp et al, 2003). But anteriorly along right MTG, the neural response to an event began to increasingly resemble its visually distinct associate and decreasingly resemble its visual matches. The influence of predictive knowledge was negatively correlated with the strength of perceptual coding along the posterior-anterior axis of right MTG, and the peak locations of these effects were reliably divergent in individual subjects.

In contrast to the divergence with perceptual coding, relational category coding increased spatially in tandem with associative coding along right MTG and peaked at the same location, leading to greater relational category information in anterior than posterior right MTG. This suggests that posterior and anterior MTG encode different information about the same events: pMTG reflects only their perceptual properties, while aMTG additionally reflects memory of their predictive relations, including the way these relations generalize across variation (here, the participating objects). pMTG lacked this information despite the fact that retrieving predictive knowledge was highly task-relevant. In summary, along right MTG, as responses become more reflective of specific predictive relations, they also become more reflective of relational categories, and less reflective of perceptual features.

These findings help resolve the two puzzles we raised earlier by showing that visual discrimination and predictive relation retrieval are handled by distinct parts of cortex, enabling the brain to accomplish both functions distinctly. Conversely, they imply that generalized representations of relational categories and knowledge of representations of specific relations both rely on common parts of right MTG, perhaps helping the brain build the former from the latter.

Other regions exhibiting associative coding (apart from right MTG) did not exhibit information about relational categories. This suggests that right MTG may have a specialized role in bringing representations of events further away from their visual attributes and closer towards generalized, relational representations. This capacity is suggestive of a key role of this region in event understanding.

### MTG as a Core Locus of Event Memory

Our findings are in line with other work implicating various parts of MTG in event memory. Among other areas, MTG exhibits correlated patterns between watching a movie clip and then recalling it later, and correlated activity *between* participants recalling the same movie (Chen et al. 2017). MTG tends to appear as part of a network of regions including the medial and lateral prefrontal cortex, angular gyrus, and precuneus, some of which appeared in our associative coding results also. These regions form part of the default mode network, thought to be critical to both prospection and memory (Buckner, Andrews-Hanna, and Schacter 2008), and related posterior-medial network, thought to be critical to encoding episodic memories in terms of relations among items and events (Ranganath and Ritchey 2012; Ranganath and Hsieh 2016). Recently, similar areas in lateral prefrontal, precuneus, and MTG were implicated in inferring the abstract structure of a set of predictive relations (Tomov, Dorfman, and Gershman 2018). However, MTG has been relatively underexplored relative to other parts of this network, and it is notable that we found it to have a unique representational signature among them.

Work on action and event knowledge offers some insight into why MTG might have had a particularly important role in our task. MTG is the most consistent area to show selective responses to retrieving action knowledge from memory (Martin et al. 1995; Kable, Lease-Spellmeyer, and Chatterjee 2002; Kable et al. 2005; Phillips et al. 2002; Perini, Caramazza, and Peelen 2014). Nearby parts of MTG show selective responses to verbs (Bedny et al. 2008; Bedny et al. 2011; Peelen, Romagno, and Caramazza 2012) and/or event nouns (Bedny, Dravida, and Saxe 2013). These responses tend to be posterior and/or superior to our aMTG ROI, but it is likely that they fall somewhere within the perception-to-memory gradient we observe.

Finally, we did not see effects of relational categories in the region of left STG reported by Frankland & Greene (2015), where they identified representations distinguishing sentences in which two objects were related in the same way (truck hitting the ball) or in the opposite way (ball hitting the truck). This could be seen as analogous to our manipulation of whether an object caused an event or reacted to the same event. However, we probed memory representations rather than the extraction of these events from sentences. It is possible that memory-based relational categories diverge from those embedded in syntax. A direct comparison of these functions would be a fascinating topic for future research.

### Neural Organization of Long-Term Associative Memory

The neural locus of long-term associative memory and its principles of organization are hardly settled. In contrast to prior work, we failed to find strong evidence of associative coding in medial temporal lobes (Schapiro, Kustner, and Turk-Browne 2012; Hindy, Ng, and Turk-Browne 2016; Reddy et al. 2015; Garvert, Dolan, and Behrens 2017; Favila, Chanales, and Kuhl 2016). This failure could be because we probed consolidated representations, as intended by the delay between learning and scanning. It is well established that the importance of the hippocampus in associative memory declines with time (Tse et al. 2007; Yamashita et al. 2009; Winocur, Moscovitch, and Sekeres 2007), suggesting that signatures of long-term, consolidated memory of predictive relations might very well be stronger elsewhere.

The neural basis of consolidated associative memory not been widely explored. There is substantial evidence of associative coding in many parts of the inferior temporal lobes, in paradigms with long training periods (Senoussi et al. 2016; Erickson and Desimone 1999; Miyashita 1988; Sakai and Miyashita 1991). However, it has been unclear whether the neural areas responsible for associative coding are those that simultaneously support visual discrimination—functions which seem to be at odds. We show a divergence between perceptual and associative functions at the large scale, and argue that long-term memory of predictive relations is represented predominantly in areas associated with other aspects of prospection, memory, and event knowledge (though not to the exclusion of others, particularly at a fine scale). The pattern we observe also broadly coincides with an increased sensitivity to *spatial* relations in more anterior vs. posterior temporal areas (Baldassano, Beck, and Fei-Fei 2017; Kaiser and Peelen 2017).

The gradient we observed from posterior to anterior MTG also coincides with observations of a similarly oriented gradients of “temporal integration windows”, that is, where correlations between subjects viewing the same movie are affected by scrambling event order at shorter vs longer timescales (Baldassano et al. 2017; Lerner et al. 2011). If anterior areas track information across a longer period of time, they could be more influenced by predictive/associative history. However, in movie stimuli, longer timescales also convey more relational content (such as interactions among actors), another reason why our findings may coincide.

## CONCLUSION

Overall, our findings help address several previous unknowns regarding long-term predictive memory. We argue that right anterior MTG is particularly specialized for representing which specific events are predictive of each other, as opposed to which are visually similar, and that it captures generalized relational similarity across distinct contexts. By pulling apart visual and relational similarity in this way, MTG plays a pivotal role in event memory and understanding.

## ACKNOWLEDGEMENTS

We would like to thank Cristina Leon, Mira Bajaj, and Jennifer Stiso for assistance with programming, data collection, and analysis. We also thank Brice Kuhl, Christopher Honey and Janice Chen for helpful discussion of these findings. This work was supported by National Institutes of Health (P30 NS45839 to the Center for Functional Neuroimaging at the University of Pennsylvania, and R01EY021717, R01DC015359, and R01DC009209 to S.L.T-S).

## SUPPLEMENTAL TABLES

**Table S1.** Pairwise *t*-values reflecting the difference in training accuracy among the event types; positive *t*-values indicate that the event in the column is more accurate than the event in the row. Df= 33; ** = *p* < .001; \**p* < .05.

|  | Event type |  |  |
| --- | --- | --- | --- |
|  | Random | Effect | Rare |
| Cause | 3.91** | 4.14** | 3.79** |
| Random |  | 2.38* | 1.82 |
| Effect |  |  | 0.43 |

**Table S2.**
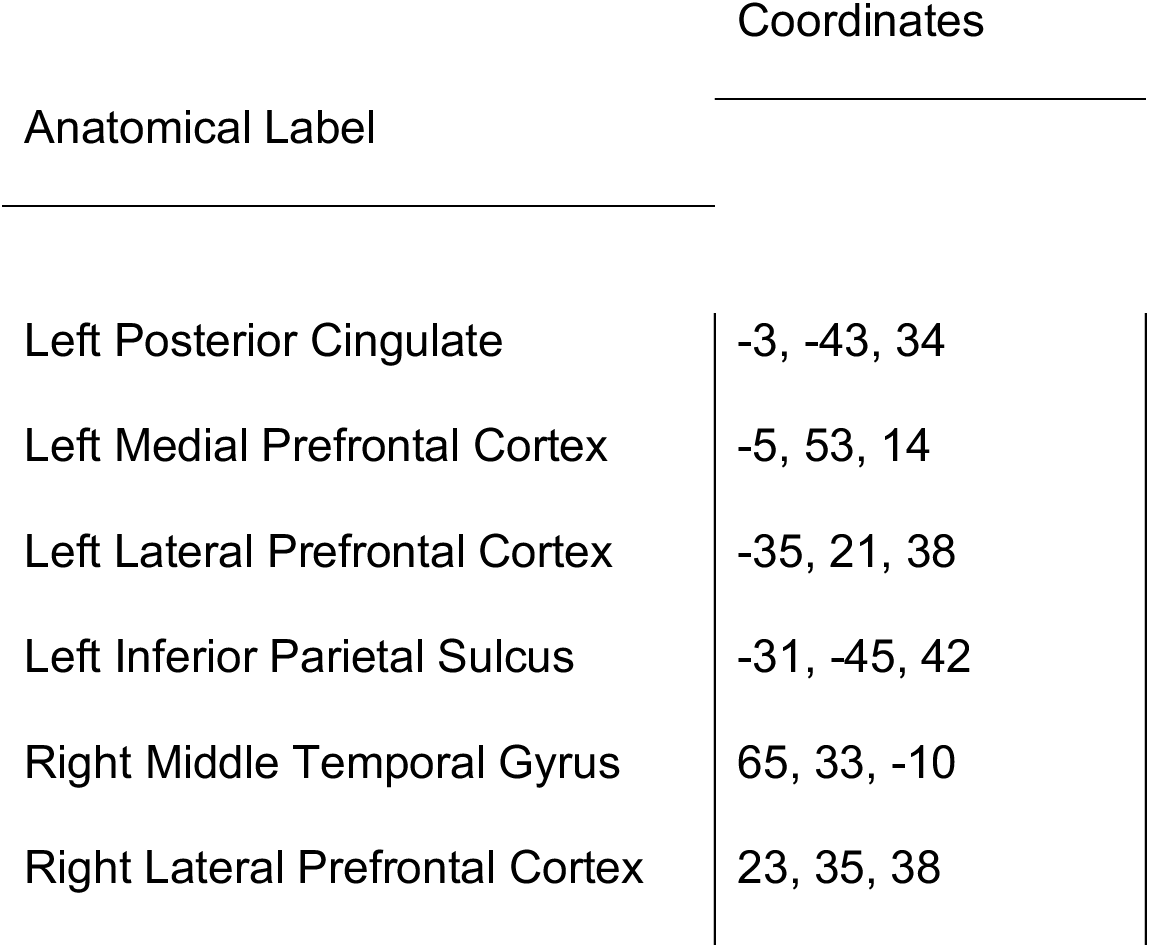
Talairach coordinates of volume-based searchlight results for the associative coding model, at their peak t-value, which correspond to significant clusters in the surface-based searchlight analysis.

**Table S3.** Mean model fits (correlation difference), standard error, *t* and *p* values for effects of perceptual model in each associative coding functional ROI. Corrected threshold for significance given 6 ROIs is *p* < .008.

| Region | Mean (SE) | $t$ | $p$ |
| --- | --- | --- | --- |
| Left Posterior Cingulate | 0.026 (0.008) | 3.22 | .001 |
| Left Medial Prefrontal Cortex | 0.015 (0.006) | 2.47 | .009 |
| Left Lateral Prefrontal Cortex | 0.027 (0.007) | 3.97 | < .001 |
| Left Inferior Parietal Sulcus | 0.042 (0.009) | 4.51 | < .001 |
| Right Middle Temporal Gyrus | 0.014 (0.007) | 1.96 | .029 |
| Right Lateral Prefrontal Cortex | 0.009 (0.006) | 1.45 | .079 |

## SUPPLEMENTAL FIGURE

**Figure S1.**
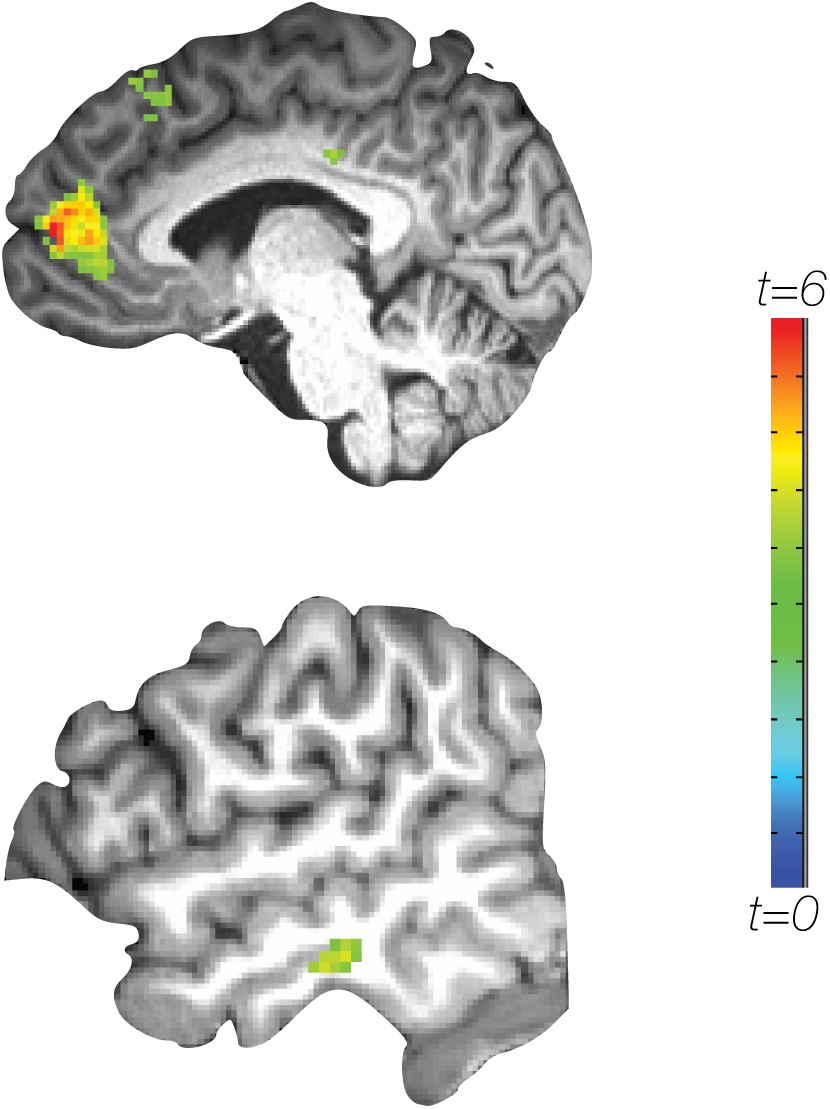
Results of volume-based searchlight for the associative coding model, thresholded at *p* < .001 uncorrected with a cluster threshold of 100 voxels (corrected threshold unknown; shown for illustration purposes only).

## Footnotes

1 Some of this work adopts the analytic approach of seeing correlated multi-voxel responses between cue and outcome stimuli, while others use classifiers trained out outcomes to test neural patterns in response to cues. We treat these as equivalent signatures.

2 In fact, we find here and elsewhere that participants see strongly predictive events like these as causally related (Leshinskaya & Thompson-Schill, under review). However, this terminology is not specifically necessary except for convenience.

3 The pre-registration had indicated the window would be 9 days, but we had one exception in which a participant was scheduled with a delay of 11 days.

